# Protein-protein docking using learned three-dimensional representations

**DOI:** 10.1101/738690

**Authors:** Georgy Derevyanko, Guillaume Lamoureux

## Abstract

Protein-protein interactions are determined by a number of hard-to-capture features related to shape complementarity, electrostatics, and hydrophobicity. These features may be intrinsic to the protein or induced by the presence of a partner. A conventional approach to protein-protein docking consists in engineering a small number of spatial features for each protein, and in minimizing the sum of their correlations with respect to the spatial arrangement of the two proteins. To generalize this approach, we introduce a deep neural network architecture that transforms the raw atomic densities of each protein into complex three-dimensional representations. Each point in the volume containing the protein is described by 48 learned features, which are correlated and combined with the features of a second protein to produce a score dependent on the relative position and orientation of the two proteins. The architecture is based on multiple layers of SE(3)-equivariant convolutional neural networks, which provide built-in rotational and translational invariance of the score with respect to the structure of the complex. The model is trained end-to-end on a set of decoy conformations generated from 851 nonredundant protein-protein complexes and is tested on data from the Protein-Protein Docking Benchmark Version 4.0.

## 1 Introduction

Proteins are the cogs of cellular machinery. Understanding how these molecules interact and predicting the structures of their complexes is an obligatory step toward understanding the emergence of the phenotype of an organism from its genotype.

The first computational prediction of the structure of a protein-protein complex was attempted by Wodak & Janin (1978). However, this prediction problem remained practically intractable until Katchalski-Katzir et al. (1992) introduced a new class of surface matching algorithms that could sample all rotations and translations of one interacting protein (the “ligand”, 𝓛) with respect to another (the “receptor”, 𝓡). The key insight of this work is that one can represent the conformational energy of the protein-protein complex as a sum over correlations of the receptor and ligand representations. It allowed sampling of the space of translations using fast Fourier transform (FFT), thus significantly reducing the complexity of the algorithm.

A similar algorithm was also used for molecular replacement in X-Ray crystallography Crowther & Blow (1967).

Since this groundbreaking work, research efforts were split into two directions: (1) accelerating rotational and translational search and (2) improving representations of proteins. The first direction saw great progress with the introduction of spherical harmonics decomposition, which allowed using fast Fourier transforms to accelerate the sampling of rotational degrees of freedom (Ritchie & Kemp, 2000). More recently, the joint rotational and translational space sampling method of Padhorny et al. (2016) further improved the execution time of rigid-body docking algorithms. The second research direction, by contrast, saw no major improvement since the introduction of electrostatic potential (Gabb et al., 1997) and the decomposition of pairwise potentials into correlation functions (Kozakov et al., 2006; Neveu et al., 2016).

In the present paper, we introduce end-to-end representation learning of protein-protein interactions that is consistent with correlation-based algorithms. Presented here as a proof of principle, this end-to-end learning approach allows future work to reap the benefits of fast conformational sampling while preserving the richness of learned representations. In Section 2 we introduce the mathematical formulation of rigid-body docking algorithms. In Section 3 we describe our model and explain how it fits within the general framework of docking algorithms. Sections 4 and 5 describe the loss function and dataset used to train the model. Finally, Section 6 is devoted to the evaluation of our approach on the Protein-Protein Docking Benchmark Version 4.0 (Hwang et al., 2010).

## 2 Rigid-body docking algorithm

Rigid-body docking algorithms usually make the assumption that there exists an energy function *E* that reaches an absolute minimum at the native position of the ligand with respect to the receptor. The form of this function constrains the algorithms we can use to efficiently sample the space of conformations. For example, energy function 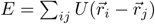, where *U* is the pairwise interaction potential between atoms *i* and *j*, requires the atomic positions to be recomputed for each ligand rotation and translation sampled.

In the present work, we express the energy function as a sum of correlations of features defined on a 3D grid. This form of *E* allows us to use well-known algorithms for accelerating sampling of translations of the ligand. For instance, let’s suppose that two interacting molecules are represented as *N* functions on a 3D grid. We denote the 3D representation of the receptor as 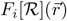 and the 3D representation of the ligand as 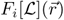, with index *i* ∈ [1, *N*]. Here, *F* denotes some functional or algorithm that computes the representations. The energy function can be computed using the following expression:

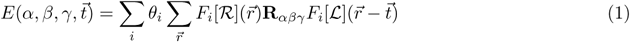

where **R**_*αβγ*_ is the rotation operator, parameterized by Euler angles (*α, β, γ*), that acts on the ligand representation. Parameters *θ*_*i*_ encode the contribution of each feature correlation into the energy of a given conformation. Expression 1 can be efficiently computed for all translations simultaneously using fast Fourier transform (FFT).

To illustrate the mechanism of the functional *F*, we construct a simple representation similar to the surface matching energy introduced by Katchalski-Katzir et al. (1992). There, *F* would output two features for any given protein 𝒫: one representing the molecular volume and another representing the molecular surface

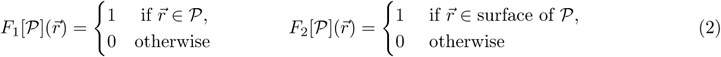

The coefficients *θ*_*i*_ should be defined in a way to penalize volume overlap and encourage surface overlap: *θ*_1_ = 1 and *θ*_2_ = − 1, for instance. In this example, the representations can be computed either from atomic densities of the molecule or from its surface mesh. Algorithm 1 below describes the workflow of the optimal rotation and translation search.

### Algorithm 1 Rigid-body docking algorithm

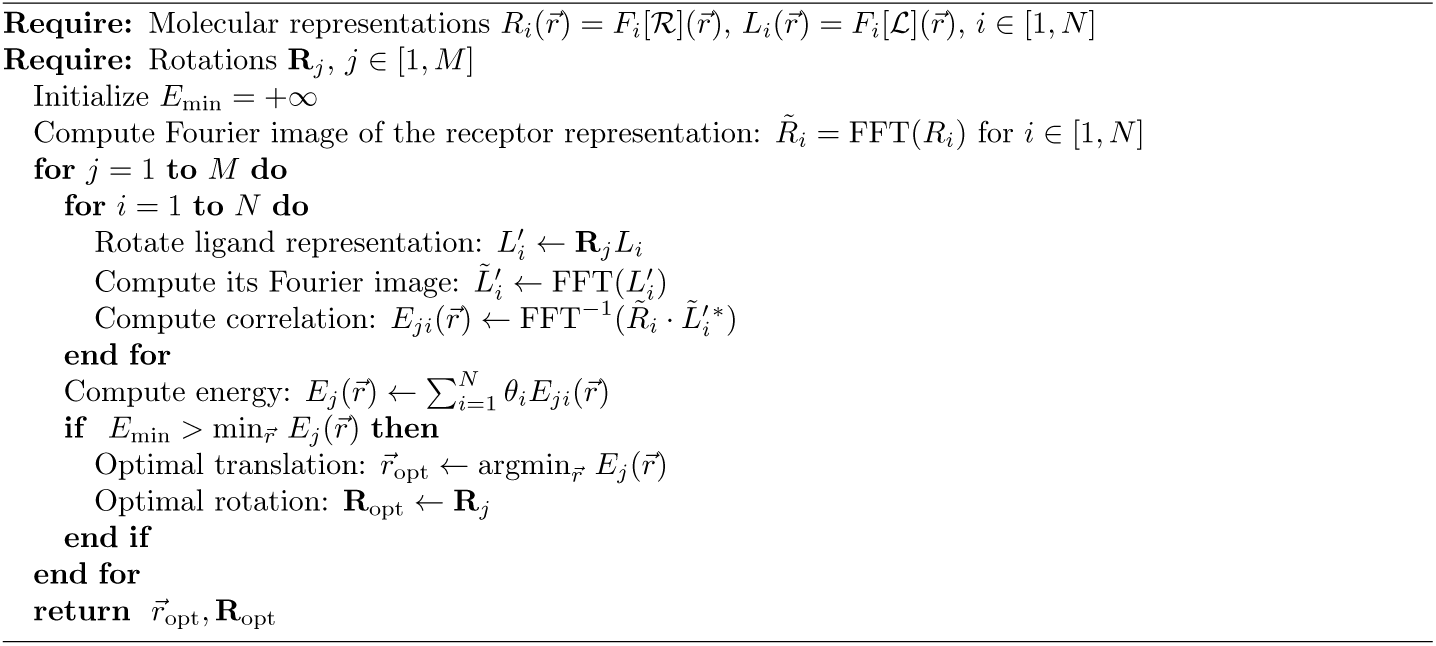

In this example, we see that the representations of the ligand and receptor are symmetric: they use the same function *F* to convert the atomic coordinates to the feature maps. According to Algorithm 1, function *F* has to be equivariant with respect to translations: 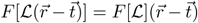. Ideally, *F* would also be equivariant with respect to rotations: 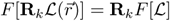. This property is implied in the description of Algorithm 1, when we compute the representation of the rotated ligand without recomputing the representation itself.

## 3 Protein representation model

The goal of this work is to demonstrate how the representations of the interacting molecules can be learned from the data. Therefore we have to construct a function *F* that converts protein atomic coordinates into features on a 3D grid, while being equivariant with respect to translations and rotations. First we project the protein atomic densities on a grid at resolution *a*^(0)^ = 1.25 Å, according to the atom types and procedure described in (Derevyanko et al., 2018, SI Section 2). Then we compute the protein representation using an SE(3)-equivariant convolutional network (Weiler et al., 2018). The diagram of the network is shown on Figure 1. The output of this model consists of two sets of volumes, 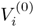 and 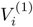, which encode information about the protein structure at two resolutions, *a*^(0)^ and *a*^(1)^ = 2*a*^(0)^. The coarsening of the resolution is the result of the strided convolution operation, which reduces each volume size by half. The projection of the atomic densities commutes with the operations from the SE(3) group, therefore the whole representation is equivariant with respect to rotations and translations of the protein. We have limited our model to isotropic filters (with *l* = 0), because training higher-order convolution filters proved to be challenging. The model architecture was mostly dictated by the memory constraints presented by the GPUs we used for training.

**Figure 1:**
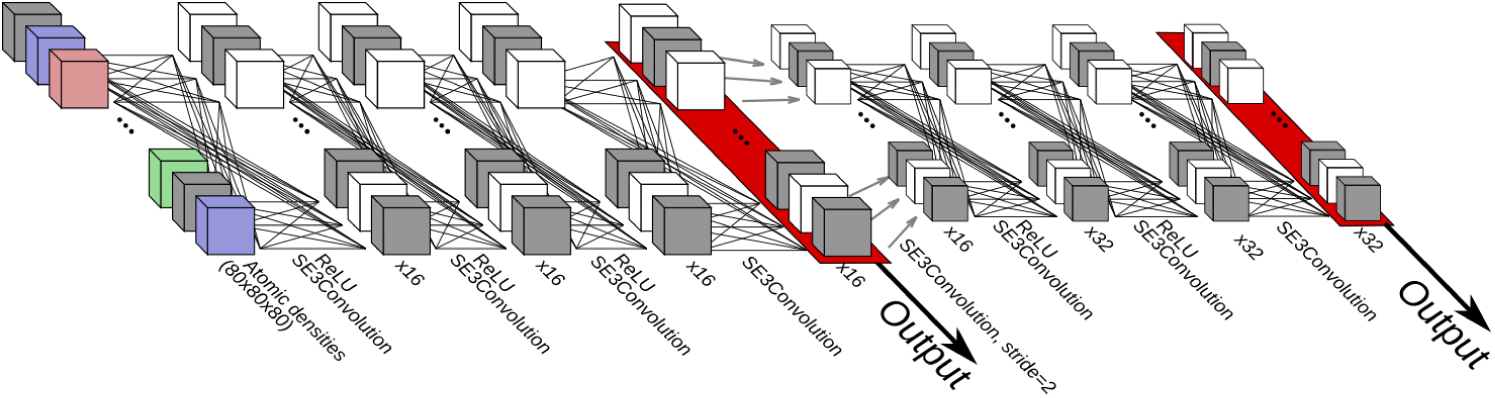
Diagram of the model that computes the representation of a protein for docking. Cubes denote 3D grids, lines connecting the cubes represent SE(3)-equivariant convolutions with spherically symmetric filters and ReLU non-linearities, and gray arrows represent strided convolutions or max pooling with stride 2. Red rectangles show the activations used as the representation of the protein, computed by the model.

The energy function of a receptor-ligand conformation is defined in the following way:

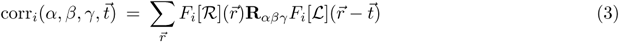

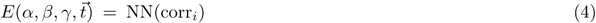

In contrast with Equation 1, we use a two-layer neural network “NN” instead of a linear combination, because it allows us to build a richer model without increasing the computational complexity.

As a baseline, we also train a standard convolutional network, which is not equivariant with respect to rotations. Detailed description of both models can be found in SI Table 1.

## 4 Loss function

Equation 3 is differentiable with respect to the parameters of functional *F*. However, there are two approaches to optimizing it: one can sample individual translations and rotations (**R**_*αβγ*_ and 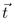) or sample all translations for a given rotation at once. In this work we explore the first approach. We generate examples of protein complexes and assign them a numerical value (the “quality”, an integer *Q* between 0 and 4) depending on how close they are to the native structures obtained experimentally. Each example corresponds to one specific rotation and translation of the ligand with respect to the receptor. The goal of the model is then to assign a score to each rotation and translation, such that it ranks them according to the quality of the complex corresponding to said rotation and translation. Figure 2 illustrates the computation of the energy for a given complex.

**Figure 2:**
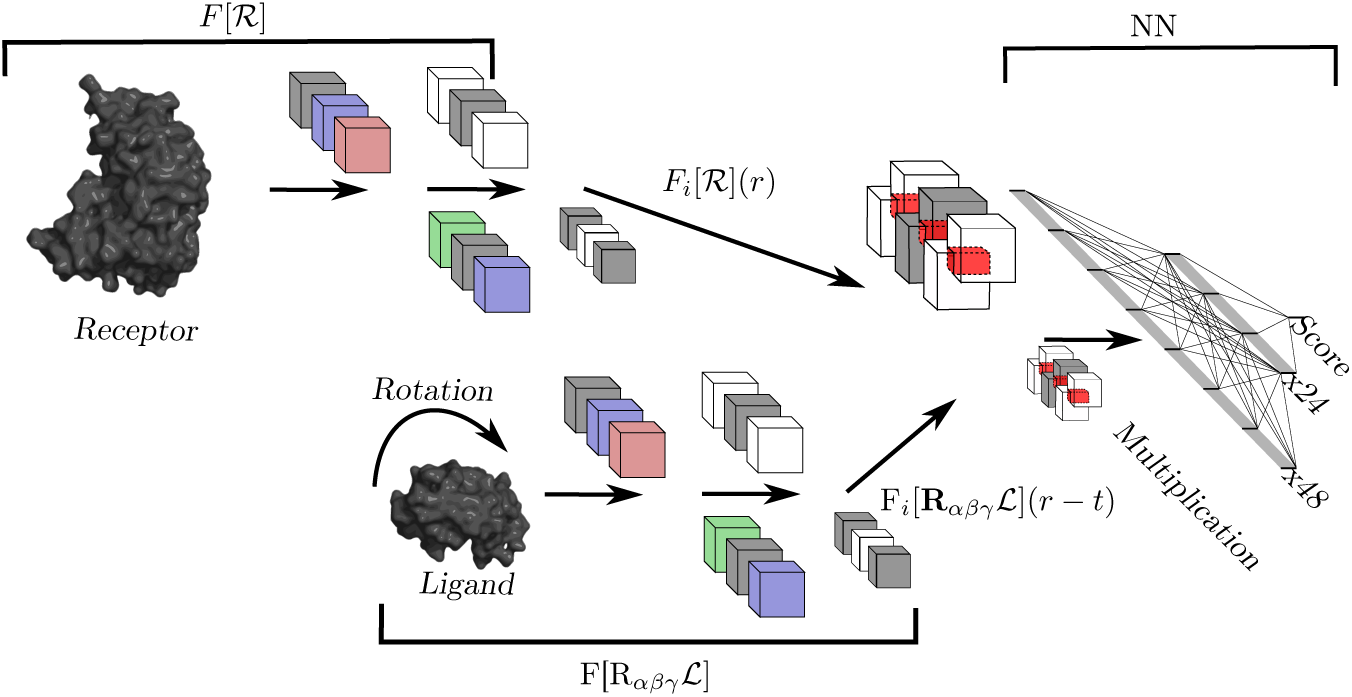
Diagram of the training process. Atomic densities of the receptor and rotated ligand are projected on 3D grids and the representations *F*_*i*_[𝓡] and *F*_*i*_[**R**_*αβγ*_𝓛] are computed. Then, translation 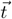 is applied to the ligand representation volumes. The two representations are multiplied and their sums are passed to a two-layer neural network to compute the energy. The red rectangles denote the overlap of the volumes after the translation is applied.

We place a number of additional conditions on the representation models (*F*) and on the neural network that ranks conformations (NN). First, the output scores produced by the model can be interpreted as an energy function that corresponds to the probability distribution over rotations and translations of the ligand with respect to the protein. The first condition on the models is dictated by the fact that all conformations where two proteins are far from each other 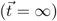 should be equally probable and should have the same score. When the bounding boxes of the two proteins do not overlap, the multiplication of their representations gives the zero vector. Therefore the first term in the loss function is:

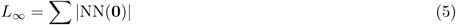

The second condition stems from the fact that, if one of the proteins is absent, the rotations and translations of the other protein are equally probable and have the same score. To make this condition consistent with the previous one, we set this score to zero. In this work we satisfy this condition by setting all the biases in the convolutional layers of the representation models to zero.

The third condition is that the generated models of the two proteins should have an energy lower than that of any conformation with 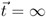. This threshold score is set to zero by the first condition. Therefore we get the following term in the loss function:

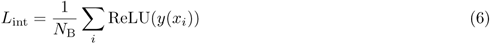

where *y*(*x*_*i*_) is the output of the model on example *x*_*i*_. *N*_B_ denotes the batch size.

Finally, the main term in the loss function is the batch ranking loss, which ranks models according to their quality. We use the margin ranking loss (Gong et al., 2013) to learn to rank for each pair of examples in a batch. Let *Q*_*i*_ denote the quality of model *i* and let *y*_*ij*_ be the ordering coefficient of models *i* and *j*, equal to +1 if *Q*_*i*_ < *Q*_*j*_ and to −1 otherwise. Let *y*_*i*_ denote the output of the model for conformation *i*. We use the following expression for the pairwise ranking loss:

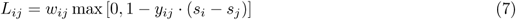

The coefficient *w*_*ij*_ is defined so that models that have the same quality are excluded from the ranking loss: *w*_*ij*_ is one if |*Q*_*i*_ − *Q*_*j*_| > 0 and is zero otherwise. For a batch of *N*_B_ examples the ranking loss is:

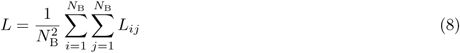

From our experiments the regularization terms go to zero during the first epochs, therefore it is safe to set all the weights to be equal: *L*_total_ = *L*_*∞*_ + *L*_int_ + *L*.

## 5 Dataset

For the sake of comparing with other algorithms that use a data-driven approach to docking (Kozakov et al., 2017, 2006; Neveu et al., 2016), we generated our datasets from the set of structures published by Huang & Zou (2008). The full set consists of 851 non-redundant protein complexes, of which 655 are homodimers and 196 are heterodimers.

To take into account the structural changes of the molecules upon binding, we generated two sets of decoys for all ligands and receptors using the NOLB method (Hoffmann & Grudinin, 2017) with RMSD cutoffs of 1.5 Å and 3.0 Å. These decoys were then docked using the Hex algorithm (Ritchie & Venkatraman, 2010), which takes only shape complementarity into account. We tuned the parameters of the Hex algorithm such that, for the majority of complexes, we obtain at least 5 ligand positions of acceptable quality. We define the quality *Q* of a decoy according to the CAPRI ranking criteria (Méndez et al., 2003) (see Supporting Information, Section 3). In total, we generated 200 decoys for each of the 851 initial structures. Complexes having less than 5 decoys of acceptable quality were excluded from the training set. We also excluded several structures that were comprised of proteins whose bounding box size was more than 80 Å. The statistics of the generated dataset are reported in Section 4 of Supporting Information.

The final dataset was split into training and validation subsets. During the training, each batch is formed by randomly sampling from the decoys of a single complex, such that it has approximately equal numbers of acceptable (*Q* ≥ 1) and incorrect (*Q* = 0) conformations.

## 6 Evaluation and Results

Models were trained using a TitanX Maxwell GPU and the evaluation was performed on four nodes with K40 GPUs (Parashar et al., 2018). On the TitanX GPU, one update takes 5 s for the CNN model (222,881 parameters) and 7 s for the SE(3)-equivariant CNN model (14,785 parameters). The details of the models architecture can be found in SI Table 1. Training loss curves on Figure 3 show that the SE(3)-equivariant representation models reach lower training loss than the ordinary model, however the validation losses are on average similar for both models. We evaluated both models after 100 epochs of training, since the validation losses had appeared to reach a minimum at this point.

**Table 1:** Performance of ordinary convolutional network (CNN) and equivariant (SE(3) CNN) representations in comparison with the ClusPro algorithm (Kozakov et al., 2006, 2017). Columns “# < 10 Å” show the average number of near-native docked structures with an interface RMSD (“IRMSD”) less than 10 Å among the 1000 lowest-scoring conformations. Columns “Min IRMSD” show the average value of the lowest interface RMSD among the 1000 lowest-scoring conformations. All averages are computed over the number of targets in the corresponding category. The difficulty of an example is defined by the interface RMSD value between bound and unbound complex. “Easy” cases have IRMSD *<* 1.5 Å and a fraction of native contacts *>* 0.4, “hard” cases have IRMSD *>* 2.2 Å, while all other cases are classified as “medium” difficulty. In the case of antibody-antigen complexes, we compared our results with ClusPro unbound and unmasked docking results.

| Enzyme-inhibitor complexes (unbound) |  |  |  |  |  |  |
| --- | --- | --- | --- | --- | --- | --- |
| Category | CNN |  | SE(3) CNN |  | ClusPro |  |
| | # $< 10$ Å | Min IRMSD | # $< 10$ Å | Min IRMSD | # $< 10$ Å | Min IRMSD |
| Easy | 543.1 | 4.24 | 480.9 | 3.97 | 172.3 | **2.94** |
| Medium | 468.0 | **5.23** | 339.5 | 6.25 | 42.5 | 8.21 |
| Hard | 97.0 | 9.12 | 197.4 | **6.87** | 19.0 | 15.9 |
| Other type complexes (unbound) |  |  |  |  |  |  |
| Easy | 189.7 | 7.87 | 192.5 | **7.09** | 78.8 | 7.54 |
| Medium | 132.8 | 7.63 | 231.5 | **5.93** | 59.9 | 7.09 |
| Hard | 231.0 | **9.15** | 94.8 | 9.61 | 49.8 | 10.48 |
| Antibody-antigen complexes (unbound) |  |  |  |  |  |  |
| - | 53.9 | 11.7 | 56.6 | 11.9 | 66.5 | **5.0** |

**Figure 3:**
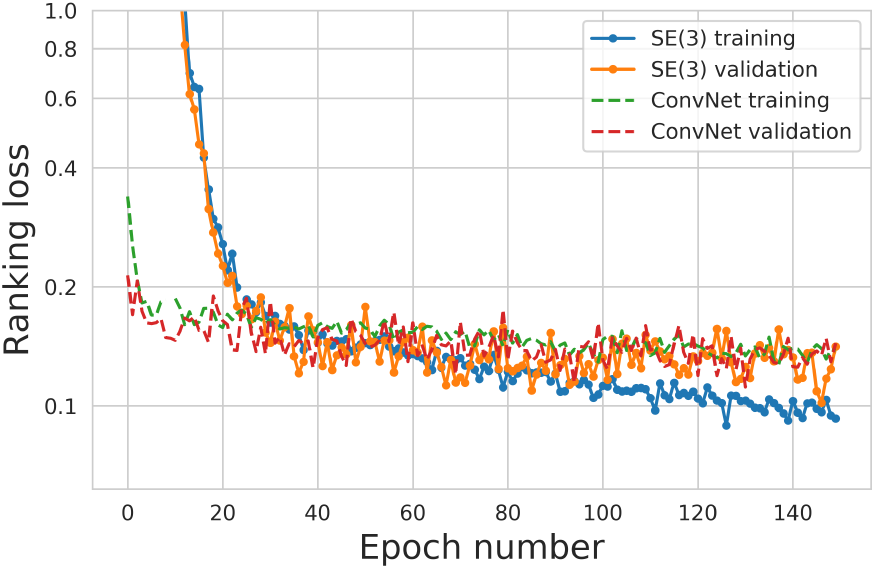
Ranking loss (Eq. 8) curves on the training and validation subsets for rotationally equivariant and ordinary representation models. Continuous lines with symbols show loss values on the validation and training sets of the equivariant model while dashed lines represent the training of the ordinary convolutional network based model. Loss values were averaged over all examples during one epoch. For clarity, the *y*-axis has a logarithmic scale. The models at epoch 100 were picked for evaluation.

During the evaluation phase we compute the correlation of the two representations for all translations simultaneously using a fast Fourier transform. Figure 4 illustrates the workflow of the evaluation phase. In the end we have 16 correlations at resolution *a*^(0)^ and 32 at resolution *a*^(1)^ (for a total of 48 spatially-resolved features). To obtain the optimal translation at the highest possible resolution, we upscale the 32 coarse correlations to resolution *a*^(0)^ and pass these volumes as features to the two-layer neural network. Finally, we get the energy in the volume at resolution *a*^(0)^. This procedure is repeated for all rotations of the ligand, covering the SO(3) group. We generate the rotation matrices using the algorithm of Yershova et al. (2010).

Figure 5 shows an example of the output score for a single rotation. The dataset does not contain any decoy in which proteins overlap (due to the method used to generate it), yet Figure 5a shows that the model ranks low all translations resulting in the two proteins overlapping. These conformations are actually excluded from the output using a “forbidden volume” feature *V*_f_, defined from *ρ*[𝓡] and *ρ*[𝓛], the atomic densities of receptor and ligand corresponding to the sums of the individual atomic densities used as input for the representation models:

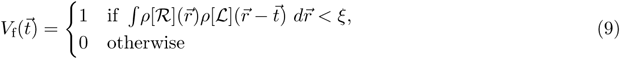

**Figure 4:**
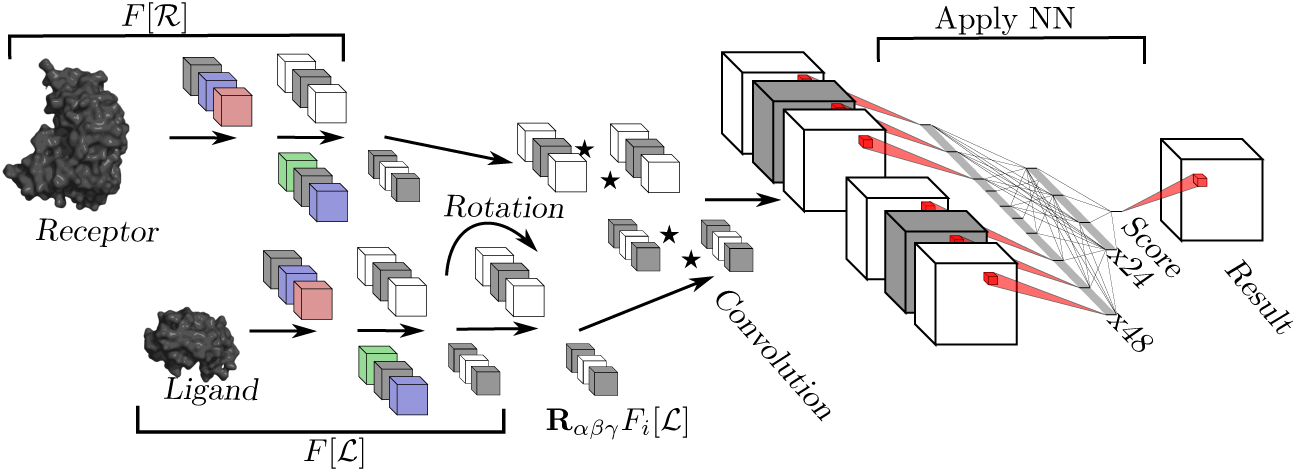
Schematic workflow of the evaluation phase of docking. Representations of ligand and receptor are computed using initial rotations and translations. Rotation is applied to the representation of the ligand, then correlations of the two representations are computed. The resulting volumes are passed through the fully-connected neural network.

**Figure 5:**
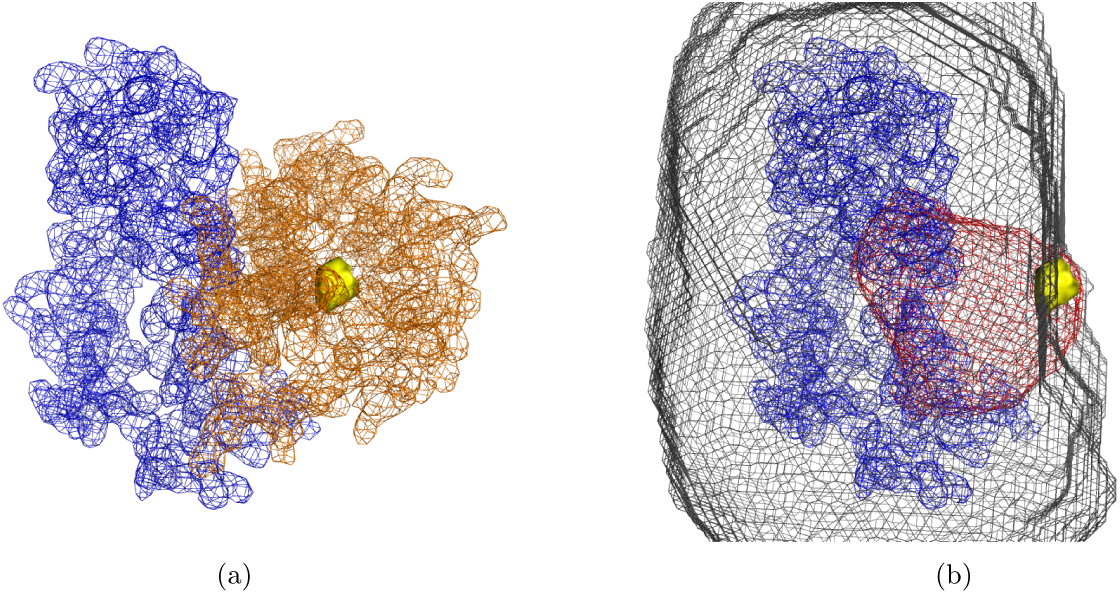
Example output of the algorithm for the homodimer PDB code 1BBH. The blue mesh shows *ρ*[𝓡], the density of the receptor (the sum of atomic densities), at isolevel 1.0. In panel (a), the orange mesh shows the density of the ligand *ρ*[𝓛] in complex with the receptor. The yellow surface shows the positions of the ligand center that generate complexes with LRMSD *<* 5.0 Å. In panel (b), the red mesh shows the output density, calculated as in Fig. 4, at isolevel −17.0. Any translation bringing the center of the ligand into that volume yields an energy of the complex below −17.0. The grey mesh shows the forbidden volume at threshold *ξ* = 300. Any translation bringing the center of the ligand into that volume creates a significant overlap with the receptor. The yellow surface, which represents the true position of the ligand, is overlapping with the red mesh and is outside the grey mesh.

The final output produced by the model is simply multiplied by *V*_f_, which means that any conformation in which the proteins overlap too much is given a zero score. The threshold *ξ* sets a limit to the overlap between the two proteins, and is a useful measure of how much flexibility we want to include in the two proteins during the docking. While preliminary experiments have shown that the forbidden volume can be learned implicitly by including additional overlapping examples to the dataset, in this work we calculate *V*_f_ with a threshold *ξ* = 300.

We evaluated our approach on the Docking Benchmark Version 4.0 (Hwang et al., 2010). To the best of our knowledge, there are only two published algorithms that use knowledge-based potentials as components of the docking procedure: ClusPro (Kozakov et al., 2006, 2017) and PEPSI (Neveu et al., 2016). In this paper we compare our approach to the ClusPro algorithm, which uses pairwise atomic potentials derived from the assumption that, in the absence of interactions, atomic distances in protein structures are distributed according to the Boltzmann distribution. The most recent benchmark of the ClusPro approach uses spherical harmonics decomposition to sample joint rotations and translations. Table 1 shows comparison between our algorithm and the results obtained using ClusPro. We evaluated our algorithm on the targets for which both the receptor and the ligand can fit within the 80 Å bounding box (corresponding to 147 of the 176 ClusPro targets). During the evaluation we used only unbound structures and we sampled the rotations with a 15° increment. ClusPro uses a finer angle increment of 5° but we found that 15° was low enough for both our CNN and SE(3) CNN models.

Taken together, our CNN or SE(3) CNN approaches outperform the state of the art in most categories, except for antibody-antigen complexes, for which ClusPro is highly accurate. This may be explained by the fact that ClusPro uses different representations for antibodies than for other targets. Our approach also appears to retain better accuracy for the more difficult cases, which display larger induced fit.

## 7 Conclusion

In this paper we describe an algorithm that enables end-to-end learning of protein representations for rigid-body docking. This general approach provides multiple benefits that remain to be explored in full. First, it allows us to include many more protein descriptors, such as electrostatics, hydrophobicity, or sequence conservation without increasing the computational complexity of the docking. In this work we used representations with 48 features, but this number can in principle be reduced to as few as 2, provided the features are rich enough. This would provide a significant speedup in evaluation without having to resort to representations based on spherical harmonics, thus avoiding problems inherent to this type of representation, such as the need to have higher-order harmonics for larger molecules. Moreover, although this work used only scalar features in the SE(3)-equivariant representation model, the approach can also be used with vector- and tensor-valued features, which are appropriate for protein properties such as solvent electrostatics and flexibility.

The method presented in this paper is directly applicable to flexible docking, since the final score of a protein complex is end-to-end differentiable, and could be optimized with respect to the atomic coordinates of each interacting protein. Finally, this work constitutes an essential building block for developing end-to-end models of protein interactomes, going from sequences to interactions.

## Acknowledgments

We thank Yoshua Bengio for hosting one of us (G.D.) at the Montreal Institute for Learning Algorithms (MILA) during some of the critical stages of the project, and for access to the MILA computational resources. This work was supported by a grant from the Natural Sciences and Engineering Research Council of Canada to G.L. (RGPIN 6658-2016). This research used resources from the Rutgers Discovery Informatics Institute (RDI2), which are supported by Rutgers and the State of New Jersey.

## Availability

The code is available at https://github.com/lupoglaz/DeepLocalProteinDocking.

