## Supporting Information for "Protein-protein docking using learned three-dimensional representations" (from another version of this preprint: version 1, 2019-08-19)

### Supplementary Information for: Protein-protein docking using learned three-dimensional representations

Georgy Derevyanko*
Department of Chemistry and Biochemistry
Concordia University
Montréal, Québec, Canada


Guillaume Lamoureux†
Department of Chemistry and Center for Computational and Integrative Biology
Rutgers University – Camden
Camden, New Jersey, USA


August 16, 2019

*https://github.com/lupoglaz
†http://lamoureuxlab.org

### 1 Models

Table 1: Details of the model architecture.

| Layer | CNN | | | | | SE(3) CNN | | | | |
|---|---|---|---|---|---|---|---|---|---|---|
| | Layer | # input | # output | Size | Stride | Layer | # input | # output | Size | Stride |
| | Input atomic densities, resolution 1.25 Å | | | | | | | | | |
| 1 | Conv3D | 11 | 16 | 5 | 1 | SE3Conv | 11 | 16 | 5 | 1 |
| 2 | ReLU | | | | | ReLU | | | | |
| 3 | Conv3D | 16 | 16 | 3 | 1 | SE3Conv | 16 | 16 | 5 | 1 |
| 4 | ReLU | | | | | ReLU | | | | |
| 5 | Conv3D | 16 | 16 | 3 | 1 | SE3Conv | 16 | 16 | 5 | 1 |
| 6 | ReLU | | | | | ReLU | | | | |
| 7 | Conv3D | 16 | 16 | 3 | 1 | SE3Conv | 16 | 16 | 5 | 1 |
| | Output #1, resolution 1.25 Å | | | | | | | | | |
| 8 | MaxPool3D | 16 | 16 | 5 | 2 | SE3Conv | 16 | 32 | 5 | 2 |
| 9 | ReLU | | | | | ReLU | | | | |
| 10 | Conv3D | 32 | 32 | 3 | 1 | SE3Conv | 32 | 32 | 5 | 1 |
| 11 | ReLU | | | | | ReLU | | | | |
| 12 | Conv3D | 32 | 32 | 3 | 1 | SE3Conv | 32 | 32 | 5 | 1 |
| 13 | ReLU | | | | | ReLU | | | | |
| 14 | Conv3D | 32 | 32 | 3 | 1 | SE3Conv | 32 | 32 | 5 | 1 |
| | Output #2, resolution 2.5 Å | | | | | | | | | |
| Total | 222,881 parameters | | | | | 14,785 parameters | | | | |

### 2 Atom types and atomic densities

Each protein structure is represented by 11 density maps corresponding to the atom types defined in Table 2. These atom types are a simplification of the 20 types proposed by Huang & Zou (2008). We used these atom types in our previous work (Derevyanko et al., 2018).

The density of an atom is represented using the function

$$\rho(r) = \begin{cases} e^{-\frac{r^2}{\sigma^2}} & \text{if } r \leq 2\sigma \text{ Å}, \\ 0 & \text{otherwise} \end{cases} \tag{1}$$

where $\sigma$ is the resolution of the grid. The atomic density is projected to the grid corresponding to its atom type. The input grids have a resolution of 1.25 Å and have $80 \times 80 \times 80$ cells.

Table 2: Atom types used in this work. Atoms in each group are identified using their standard PDB residue names and atom names. Asterisks (*) correspond to either 1, 2, or 3. Table reproduced from Derevyanko et al. (2018).

| Type | Description | Atoms |
|---|---|---|
| 1 | Sulfur/selenium | CYS:SG, MET:SD, MSE:SE |
| 2 | Nitrogen (amide) | ASN:ND2, GLN:NE2, backbone N (including N-terminal) |
| 3 | Nitrogen (aromatic) | HIS:ND1/NE1, TRP:NE1 |
| 4 | Nitrogen (guanidinium) | ARG:NE/NH* |
| 5 | Nitrogen (ammonium) | LYS:NZ |
| 6 | Oxygen (carbonyl) | ASN:OD1, GLN:OE1, backbone O (except C-terminal) |
| 7 | Oxygen (hydroxyl) | SER:OG, THR:OG1, TYR:OH |
| 8 | Oxygen (carboxyl) | ASP:OD*, GLU:OE*, C-terminal O, C-terminal OXT |
| 9 | Carbon (sp2) | ARG:CZ, ASN:CG, ASP:CG, GLN:CD, GLU:CD, backbone C |
| 10 | Carbon (aromatic) | HIS:CG/CD2/CE1, PHE:CG/CD*/CE*/CZ, TRP:CG/CD*/CE*/CZ*/CH2, TYR:CG/CD*/CE*/CZ |
| 11 | Carbon (sp3) | ALA:CB, ARG:CB/CG/CD, ASN:CB, ASP:CB, CYS:CB, GLN:CB/CG, GLU:CB/CG, HIS:CB, ILE:CB/CG*/CD1, LEU:CB/CG/CD*, LYS:CB/CG/CD/CE, MET:CB/CG/CE, MSE:CB/CG/CE, PHE:CB, PRO:CB/CG/CD, SER:CB, THR:CB/CG2, TRP:CB, TYR:CB, VAL:CB/CG*, backbone CA |

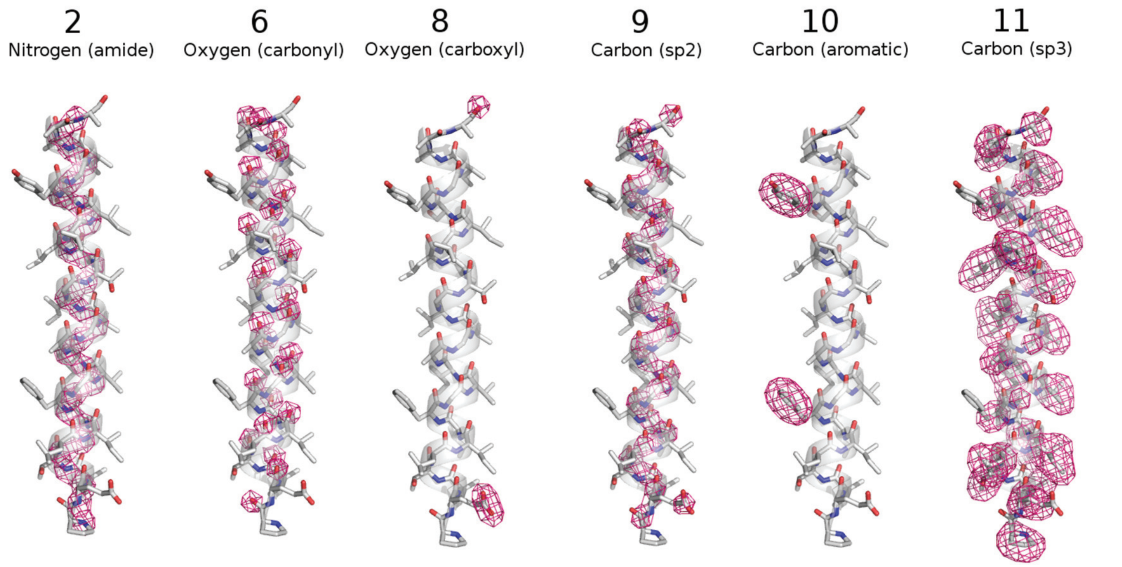


Figure 1: Atomic density representation of a protein structure (PDB code 5eh6; simple $\alpha$-helical peptide). The density maps are calculated according to Eq. 1 and rendered with an isosurface level of 0.5. The structure contains only 7 of the 11 atom types, and only those density maps are shown. (The 4 other maps have zero density everywhere.) Figure reproduced from Derevyanko et al. (2018).

#### 3 CAPRI model quality criteria

Two residues are considered in contact when any of their non-hydrogen atoms are within 5 Å. Denoting the set of contacts in the native structure as $\mathcal{C}_{\text{nat}}$ and the set of contacts in the model as $\mathcal{C}_{\text{mdl}}$, the fraction of native contacts reads:

$$f_{\text{nat}} = \frac{|\mathcal{C}_{\text{nat}} \cap \mathcal{C}_{\text{mdl}}|}{|\mathcal{C}_{\text{nat}}|} \tag{2}$$

The "ligand RMSD" (LRMSD) is defined as the root mean square deviation of the ligand in the native complex from the ligand in the model, provided the receptors in both complexes are superimposed. The root mean square deviation is calculated using the atoms of the backbone (N, $\text{C}_\alpha$, C, O) only. We denote the coordinates of the backbone atoms of the model ligand as $\mathbf{x}^{\text{lig}}_{\text{mdl}}$, and those of the model receptor as $\mathbf{x}^{\text{rec}}_{\text{mdl}}$. The corresponding atomic coordinates in the native structure are $\mathbf{x}^{\text{lig}}_{\text{nat}}$ and $\mathbf{x}^{\text{rec}}_{\text{nat}}$. The ligand RMSD reads:

$$\text{LRMSD} = \sqrt{\frac{1}{N}\sum \left|\mathbf{x}^{\text{lig}}_{\text{nat}} - \mathbf{A}\cdot\mathbf{x}^{\text{lig}}_{\text{mdl}}\right|} \quad \text{with} \quad \mathbf{A} = \underset{\mathbf{A}}{\text{argmin}}\sqrt{\frac{1}{N}\sum \left|\mathbf{x}^{\text{rec}}_{\text{nat}} - \mathbf{A}\cdot\mathbf{x}^{\text{rec}}_{\text{mdl}}\right|} \tag{3}$$

Here, $\mathbf{A}$ corresponds to a rigid transformation of the coordinates.

Interfacial residues are defined as the receptor residues that have at least one atom within 10 Å of a ligand atom, and vice versa. The "interface RMSD" (IRMSD) is defined as the root mean square deviation between the backbone atoms of the interface residues in the native structure, $\mathbf{x}^{\text{int}}_{\text{nat}}$, and the equivalent atoms in the model structure, $\mathbf{x}^{\text{int}}_{\text{mdl}}$:

$$\text{IRMSD} = \min_{\mathbf{A}}\sqrt{\frac{1}{N}\sum \left|\mathbf{x}^{\text{int}}_{\text{nat}} - \mathbf{A}\cdot\mathbf{x}^{\text{int}}_{\text{mdl}}\right|} \tag{4}$$

Table 3: Quality criteria for the models of protein-protein complexes used in CAPRI competition. Table adapted from Méndez et al. (2003) We use the rank number as an input to the batch ranking loss function.

| Rank, Rank number | $f_{\text{nat}}$ | LRMSD (Å) | IRMSD (Å) |
|---|---|---|---|
| High, 3 | $x \geq 0.5$ | $x \leq 1.0$ | or $x \leq 1.0$ |
| Medium, 2 | $x \geq 0.3$ | $1.0 < x \leq 5.0$ | or $1.0 < x \leq 2.0$ |
| Acceptable, 1 | $x \geq 0.1$ | $5.0 < x \leq 10.0$ | or $2.0 < x \leq 4.0$ |
| Incorrect, 0 | $x < 0.1$ | | |

### 4 Dataset

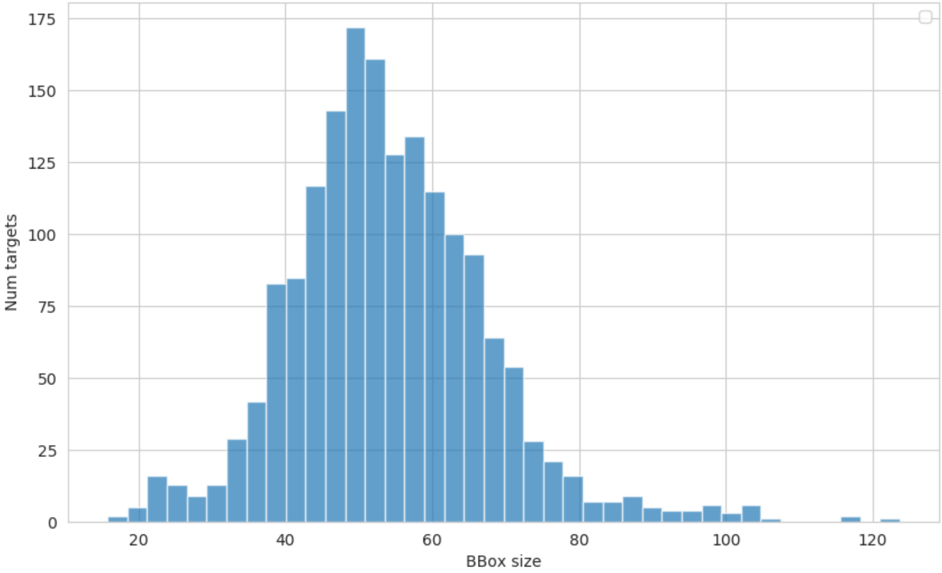


Figure 2: Distribution of the bounding box sizes for all proteins (ligands or receptors) of all 851 targets.

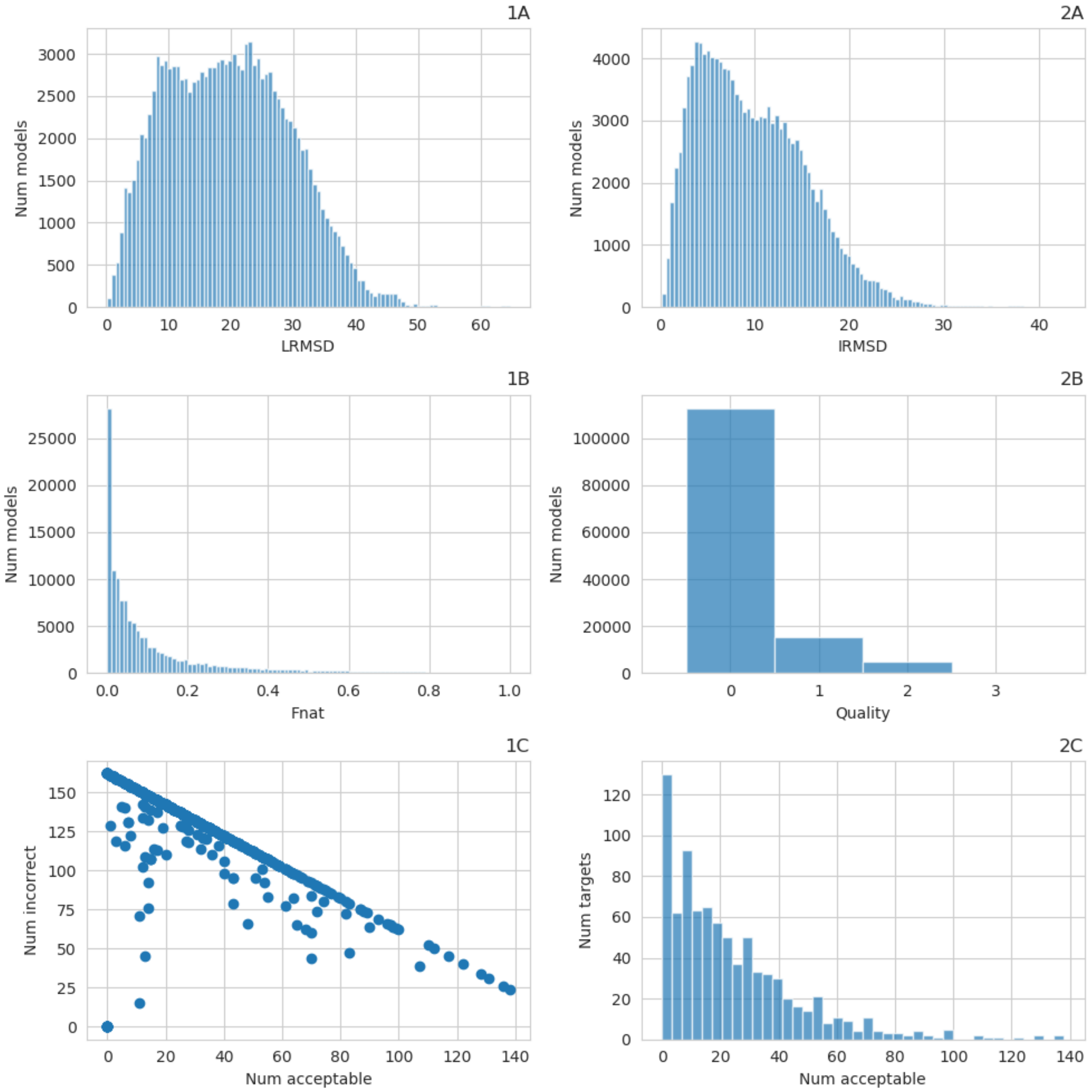


Figure 3: Distributions of LRMSD (panel 1A), IRMSD (panel 2A) and $f_{\text{nat}}$ (panel 1B). Panel 2B shows the distribution of qualities among the generated models, with 0 corresponding to an incorrect model and 3 to a high quality model, as described in Table 3 above. Panel 1C shows the number of generated models of acceptable quality (or better) versus incorrect quality for each of the 851 original targets. Panel 2C shows the distribution of targets by the number of acceptable (or better) models generated.
